# *Trypanosoma cruzi* persisters that survive benznidazole treatment *in vitro* and *in vivo* are in a transient non-replicative state

**DOI:** 10.1101/2023.08.23.554400

**Authors:** Shiromani Jayawardhana, Alexander I. Ward, Amanda F. Francisco, Michael D. Lewis, Martin C. Taylor, John M. Kelly, Francisco Olmo

## Abstract

Benznidazole is the front-line drug used to treat infections with *Trypanosoma cruzi*, the causative agent of Chagas disease. However, for reasons that are unknown, treatment failures are common. To assess the nature of parasites that persist after treatment, we first exposed infected mammalian cell monolayers to a benznidazole regimen that reduces the intracellular amastigote population to <1% of the pre-treatment level. Of host cells that remained infected, the vast majority contained only one or two surviving intracellular amastigotes. Analysis, using incorporation of the thymidine analogue EdU, revealed these surviving parasites to be in a transient non-replicative state. Furthermore, treatment with benznidazole led to widespread damage to parasite DNA. When parasites that survived treatment in mice were examined using *in vivo* and *ex vivo* bioluminescence imaging, we found that recrudescence is not due to persistence of parasites in a specific organ or tissue that preferentially protects them from drug activity. Surviving parasites were widely distributed and located in host cells where the vast majority contained only one or two amastigotes. Therefore, infection relapse does not arise from a small number of intact large nests. Rather, persisters are either survivors of intracellular populations where co-located parasites have been killed, or amastigotes in single/low-level infected cells exist in a state where they are less susceptible to benznidazole. Assessment by EdU incorporation revealed that the small number of parasites which persist in mice after treatment are initially non-replicative. A possible explanation could be that triggering of the *T. cruzi* DNA damage response pathway by the activity of benznidazole metabolites results in exit from the cell cycle as parasites attempt DNA repair, and that metabolic changes associated with non-proliferation act to reduce drug susceptibility. Alternatively, a small percentage of the parasite population may pre-exist in this non-replicative state prior to treatment.

**Author Summary:** *Trypanosoma cruzi* is the causative agent of Chagas disease, the most important parasitic infection in Latin America. For reasons that are not established, the front-line drug benznidazole often fails to achieve sterile cure. Here, we used highly sensitive imaging technology to investigate the impact of benznidazole on *T. cruzi* infected mice. Following non-curative treatment, we found that persistence is not restricted to a specific organ or tissue that preferentially protects the parasite from drug activity. Rather, surviving parasites are widely distributed, although overall tissue levels are extremely low. These persisters are located in host cells that typically contain only one or two non-replicating intracellular amastigotes. However, these parasites re-initiate DNA replication within several days of treatment cessation and begin to proliferate. Therefore, being in a non-replicative state seems to confer protection against drug-mediated trypanocidal activity. Benznidazole treatment results in widespread damage to parasite DNA. One possibility therefore, is that this triggers the *T. cruzi* DNA damage response pathway, resulting in exit from the cell cycle as parasites attempt DNA repair. Alternatively, persisters may be derived from a small parasite sub-population that pre-exists in a non-replicative state prior to treatment.

## Introduction

Chagas disease is caused by the insect-transmitted protozoan parasite *Trypanosoma cruzi* and is a major public health problem throughout Latin America, with 6 - 7 million people infected [1]. In addition, many cases are now being detected within migrant populations world-wide, particularly in Europe and the USA [2,3]. *T. cruzi* is an obligate intracellular parasite, with a wide host cell range. During the acute stage of the disease, which in humans occurs 2 - 8 weeks post-infection, parasites become widely disseminated in blood and tissues, and the infection manifests as a transient, typically mild, febrile condition. In children, the acute stage can be more serious, sometimes resulting in myocarditis or meningoencephalitis, with fatal outcomes in 5% of diagnosed cases. The acute stage is normally controlled by adaptive immune responses, mediated by CD8^+^ IFN- ^+^ T cells [4], with the infection then advancing to an asymptomatic chronic stage, in which the parasite burden is extremely low and focally restricted. However, 30 - 40% of those that are chronically infected eventually progress to a symptomatic stage, although this can take decades. Most develop cardiomyopathy, or less commonly, digestive tract megasyndromes, or both [5,6]. Infection with *T. cruzi* is a major cause of heart disease throughout endemic regions.

The nitroheterocyclic compound benznidazole is the front-line drug for *T. cruzi* infection [7,8]. Although it has been in use for almost 50 years, treatment failures are common [9–11]. Several factors have been implicated, including the effects of drug toxicity and the long administration period (60 - 90 days) on patient compliance, and the diverse nature of the *T. cruzi* species, which exhibits significant natural variation in drug susceptibility, both within and between lineages [12,13]. In addition, spontaneous parasite dormancy [14], stress-induced cell cycle arrest [15], and a reduced proliferation rate during the chronic stage [16] have all been proposed as potential mechanisms that could protect the parasite from drug treatment. Benznidazole is a pro-drug and must be activated within the parasite by the mitochondrial nitroreductase TcNTR-1 [7,12,17]. Although the precise mode of action remains to be resolved, current evidence supports a mechanism whereby highly mutagenic benznidazole metabolites, including glyoxal, cause widespread damage to genomic DNA [18–20]. Furthermore, there is a potential for cross-resistance to the other anti-*T. cruzi* drug nifurtimox, which also requires TcNTR-1-mediated activation [17,21].

In recent clinical trials, benznidazole treatment failure has been reported in 20-50% of patients [10]. Investigating the reasons for this has been complicated by the extremely low parasite burden, the focal nature of chronic infections, and the resultant difficulties in establishing cure/non-cure. It is unclear, for example, whether *T. cruzi* is able to persist in specific tissues or organs that are less accessible to benznidazole, or if a sub-set of dormant or metabolically quiescent parasites are able to survive drug exposure that kills the majority of the parasite population. To better investigate treatment failure, we have exploited a genetically modified parasite cell line that expresses a reporter fusion protein that is both bioluminescent and fluorescent [22]. The red-shifted bioluminescent component of this protein allows the tissue-specific location of parasites to be resolved with exquisite sensitivity during murine infections [23], and the fluorescent component then enables parasites to be visualized at the level of individual infected cells [24,25]. This has enabled us to localize and characterize parasites that persist after non-curative benznidazole treatment in murine models of acute Chagas disease. Our results indicate that parasites which survive treatment are in a non-replicative state.

## Results

### T. cruzi persisters that survive benznidazole treatment in vitro are in a transient non-replicative state

To generate persisters *in vitro*, we adapted the “washout” protocol described by MacLean *et al.* [26], using MA104 cells infected with the *T. cruzi* CL Luc::mNeon clone [22] (Materials and Methods). A treatment period of 8 days and a benznidazole concentration of 20 µM (10x amastigote EC_50_) was assessed as being optimal for this parasite strain:host-cell type combination (Figure 1). Under these conditions, all parasites stopped replicating within 4 days of treatment initiation (based on incorporation of the thymidine analogue EdU) (Figure 1a and b), and the amastigote population fell to <1% of the pre-treatment level. In addition, by day 5 and beyond, the majority of infected cells contained only one intracellular amastigote (Figure 1c). When benznidazole was removed, flagellated trypomastigotes did eventually develop and undergo egress, indicating the long-term viability of at least some of the surviving parasites, even after 8 days exposure to 200 µM (Figure 1d).

**Figure 1.**
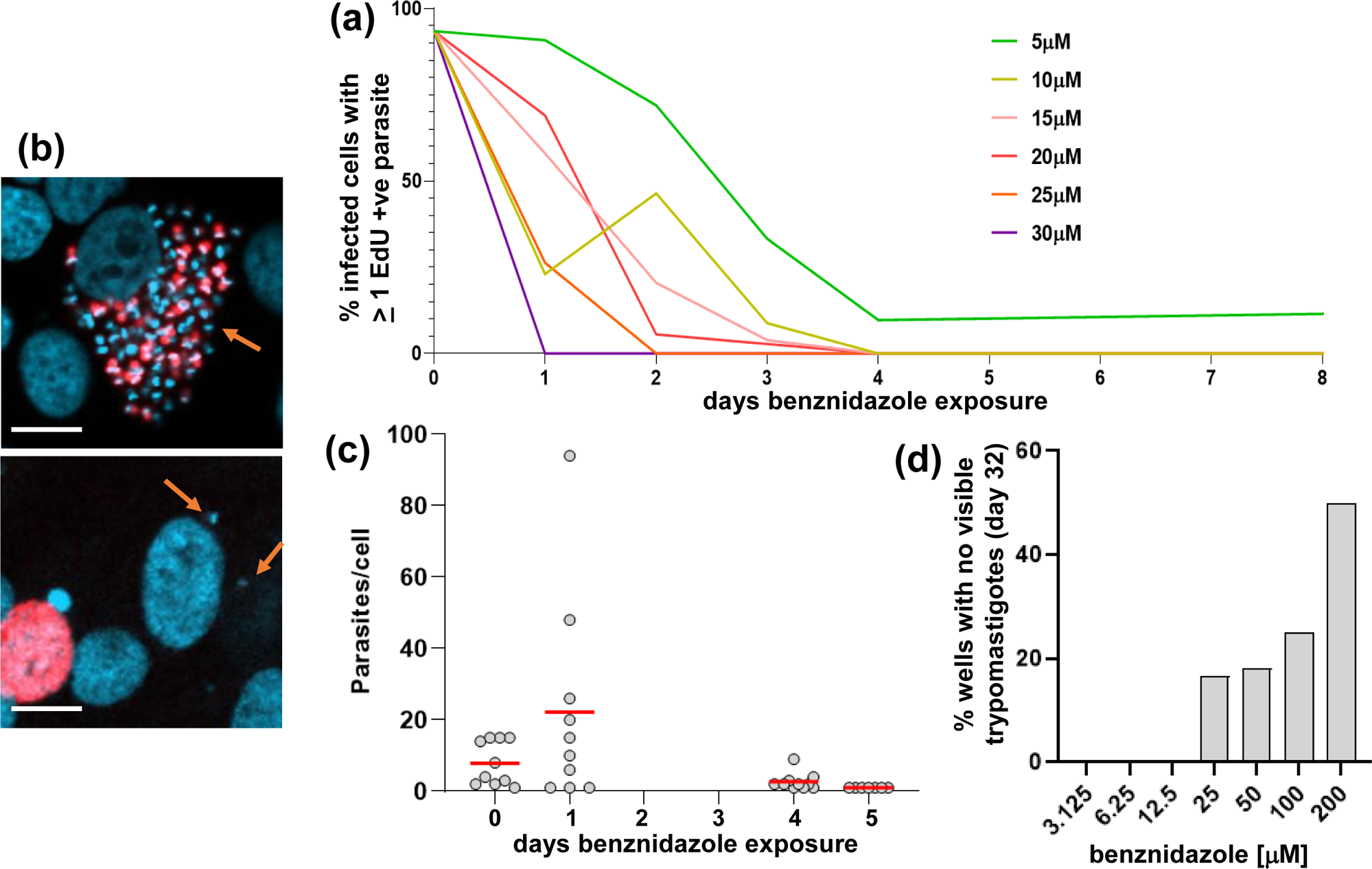
Assessing benznidazole treatment conditions to optimise the generation of *T. cruzi* persisters. (a) MA104 cells were infected with *T. cruzi* CL Luc::mNeon [22] in 24-well plates containing 13-mm diameter glass coverslips, incubated for 3 days, and then treated with benznidazole for 8 days at a range of concentrations (5-30 µM) (Materials and Methods). Each day, selected cultures were exposed to 10 µM EdU for 6 hours, fixed on coverslips, developed and scanned by fluorescence microscopy. The data are presented as % infected host cells containing at least one EdU+ve amastigote. (b) Upper image; an infected cell prior to drug exposure, with parasites that are in S-phase during the exposure period shown in red (EdU incorporation). DAPI staining (blue) identifies host cell nuclei and amastigotes, which are recognisable by their distinctive disc-like kinetoplast genome (orange arrow). Lower image; an infected cell after 8 days exposure to 10 µM benznidazole. The two amastigotes highlighted by arrows did not undergo DNA replication during the period of EdU exposure. The adjacent red stained host cell nucleus serves as a positive control for EdU labelling. White scale bars = 10 µm. (c) Infected cell monolayers in 24-well plates (as above) were treated with 20 µM benznidazole. On the days indicated, a coverslip was fixed and the amastigote content of randomly selected infected cells recorded. By day 5, all imaged infected cells contained a single amastigote. Average intracellular burden indicated by red line. (d) Infected monolayers in 24-well plates were treated with benznidazole for 8 days at the concentrations indicated. After washing, cultures were maintained in MEM for 32 days and monitored for the appearance of extracellular differentiated trypomastigotes. This assay period (a total of ∼50 days) is the limit attainable with monolayers of MA104 cells.

We further examined parasite persisters using live sorting of infected MA104 cells. Cultures of highly infected cells (Figure 1b, as example) were treated with 20 µM benznidazole for 8 days, as above, and the cells then detached to generate a suspension (Figure 2a and b, Materials and Methods). Aliquots of the suspension were incubated with propidium iodide and separated using an Aria BD Cell Sorter to determine the level of host cell viability within the population. There was a minimal level of cell death (Figure 2b and c). When host cells were separated based on the presence/absence of parasite-expressed green fluorescence, the vast majority (>99%) were found to be fluorescence-negative. The only fluorescence-positive cells detected were at the lowest gating, with each cell containing only 1, 2 or occasionally 3 parasites (Figure 2d and e). Subsequent plating of these infected cells confirmed their viability (Figure 2f), with trypomastigote egress detectable after 7-14 days in culture. Collectively, these experiments indicate that persister parasites that survive benznidazole treatment must either originate from infected host cells in which other co-habiting parasites have been eliminated, despite being exposed to equivalent drug pressure, or there is an intrinsic feature of amastigotes in single/low-level infected cells that can confer protection against benznidazole.

**Figure 2.**
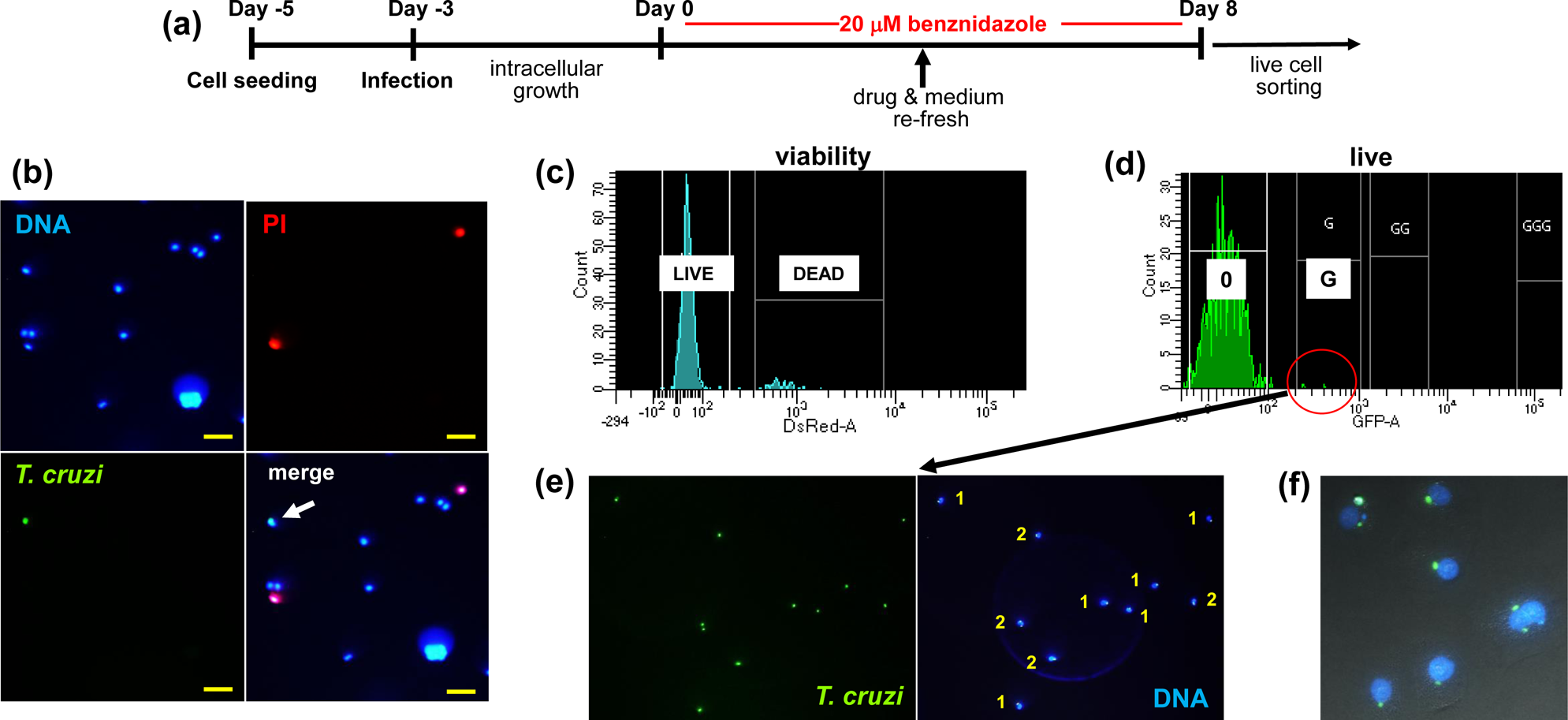
Isolation of intracellular *T. cruzi* persisters by live cell sorting following *in vitro* benznidazole treatment. (a) Experimental outline. Cultures of MA104 cells were infected with *T. cruzi* CL Luc::mNeon parasites. After 72 hours, they were treated with 20 µM benznidazole for a further 8 days. Cellular suspensions were then generated for analysis by live cell sorting (Materials and Methods). (b) Live fluorescence images of a cellular suspension showing DNA (Hoechst staining, blue), non-viable cells (propidium iodide (PI) staining, red) and parasite-infected cells (green fluorescence). In the merged image (right), the white arrow indicates a *T. cruzi* infected MA104 cell. Yellow scale bar = 50 µm. (c) Fractionation of infected cell suspension, following PI staining, using an Aria BD Cell Sorter. The small percentage of non-viable cells can be separated on the basis of acquired PI fluorescence. (d) Fractionation of live host cells into infected (within red circle) and non-infected sub-populations based on the green fluorescence of parasite persisters. 0, background fluorescence; G, lowest green fluorescence gating. (e) Image of an infected cell subpopulation suspension post-sorting. The number of parasites in each cell is indicated (1 or 2). (f) Sorted infected cells 18 hours post-plating (Materials and Methods). DNA, (blue); amastigotes, (green).

To assess the replicative status of intracellular amastigotes that persist after 8 days benznidazole exposure, we monitored the incorporation of EdU into parasite DNA (Figure 3). At selected time points post-treatment, cell monolayers were exposed to EdU for 6 hours, coverslips were removed from the 24-well plates, processed (Materials and Methods), and then scanned exhaustively to locate surviving parasites (green fluorescence). The vast majority of the remaining infected cells contained only 1 or 2 amastigotes (Figure 3a). Parasites were then further assessed to identify those that were in S-phase during the 6 hour EdU exposure period (indicated by red fluorescence). In the case of non-treated infected cells, 30-50% of intracellular amastigotes are EdU+ve under these conditions, indicative of DNA replication (Figure 1a and 3c). In contrast, of 607 intracellular amastigotes that were detected in the 9 days following cessation of drug treatment, 99.5% were EdU-ve (Figure 3d and e). However, by day 11 some surviving parasites had re-entered the cell cycle as evidenced by increased intracellular numbers and EdU positivity (Figure 3a and e). Therefore, *T. cruzi* amastigotes that persist after benznidazole treatment *in vitro* are in a non-replicative state, but retain a proliferative capacity.

**Figure 3.**
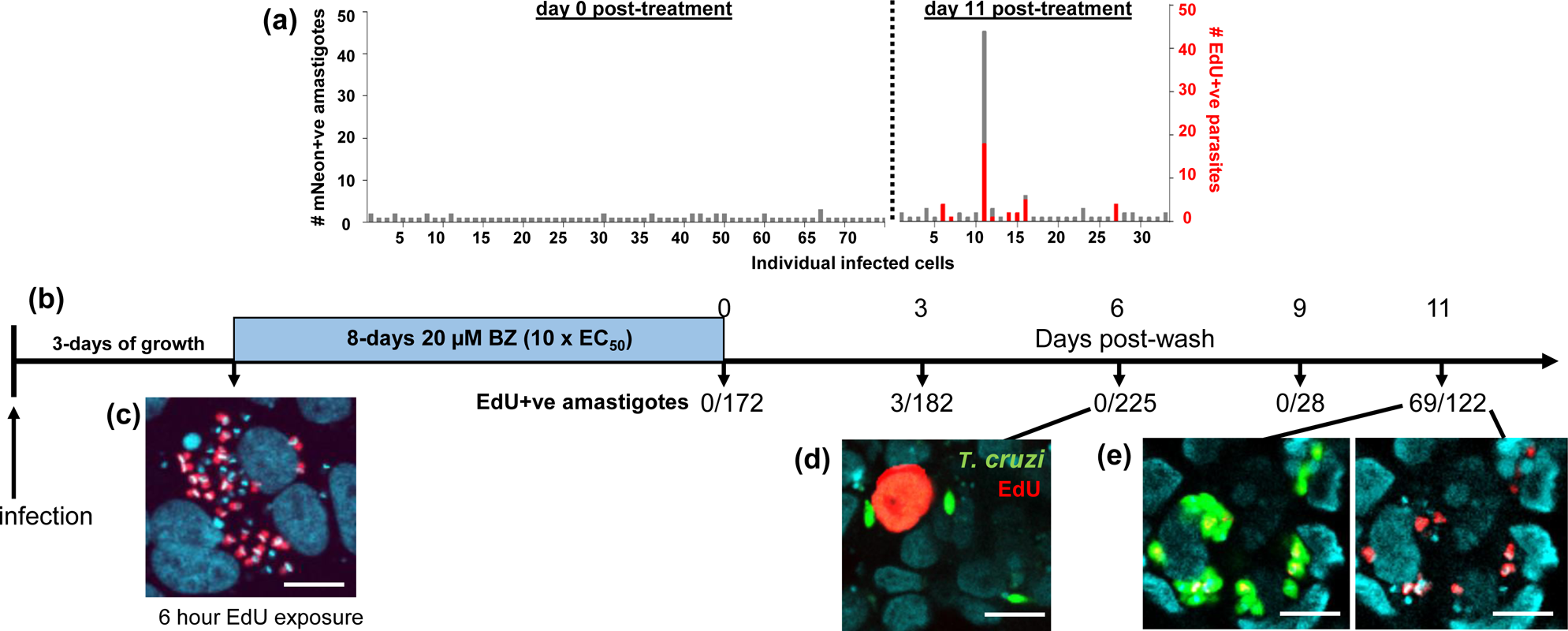
Assessing the replicative status of persister parasites *in vitro* following benznidazole treatment. MA104 cells were infected with *T. cruzi* CL Luc::mNeon in 24-well plates as indicated, and after 3 days growth, they were treated with 20 µM benznidazole for 8 days. When benznidazole was removed, the cells were maintained in complete MEM and monitored by fluorescence microscopy. To identify parasites undergoing DNA replication, cultures were exposed to 10 µM EdU for 6 hours and the coverslips processed for analysis using a Zeiss LSM880 confocal microscope (Materials and Methods). (a) Amastigote numbers in infected cells immediately after drug removal, and at day 11 post-treatment. The number of EdU+ve parasites is shown in red. (b) Timeline of an independent experiment in which the number of EdU+ve amastigotes was assessed periodically after the cessation of treatment. (c) A pre-treatment parasite nest in which ∼50% of the amastigotes are in S-phase during the period of EdU exposure (red). DAPI staining (blue) identifies host cell nuclei (large) and the parasite kinetoplast DNA (small, intense blue discs). (d) At 6 days post-wash, amastigotes (green) are in a non-replicative state. An MA104 cell in S-phase is identified by nuclear EdU staining (red). (e) Images showing parasites that have re-entered the cell cycle (day 11) and are undergoing asynchronous DNA replication [24]. White scale bars = 10 µm.

The bioactivation of benznidazole is initiated by the parasite-specific nitroreductase TcNTR-1, leading to the generation of reactive metabolites that have mutagenic properties [7,12,17–20]. A possible outcome of this could be induction of the DNA damage response system in *T. cruzi*, resulting in exit from the cell cycle and entry into a non-proliferative state [27–30]. To investigate the impact of benznidazole on the structural integrity of parasite genomic DNA, we used the TUNEL (terminal deoxynucleotidyl transferase dUTP nick end labelling) assay (Figure 4), a procedure originally developed to monitor apoptotic cell death [31]. In *T. cruzi*, this technique can be used to identify parasites undergoing replication of mitochondrial DNA (kDNA) [22,24]. Parasites early in kDNA S-phase exhibit TUNEL positivity in antipodal sites, either side of the kDNA disk, indicative of the two replication factories (see inset Figure 4b). Later in the cycle, the whole disk becomes labelled. Under normal growth conditions however, *T. cruzi* nuclear DNA does not display a detectable positive signal using this technique. As inferred from TUNEL labelling of free 3’-hydroxyl groups, 24 hours benznidazole treatment (200 µM) resulted in widespread fragmentation of parasite DNA, while the nuclei of adjacent mammalian cells remained unlabelled (Figure 4b). Under these conditions, an 8-fold extension of the treatment period is insufficient to completely eliminate parasites from infected cultures (Figure 1d). As a control, incubation with the hydroperoxide TBHP, which induces apoptosis and necroptosis [32], resulted in lesions to DNA in both parasite and host cells, with a labelling profile similar to that generated by post-fixation DNase treatment (Figure 4b). Thus, DNA damage resulting from benznidazole treatment is parasite-specific, reflecting selective metabolic reduction of the drug by TcNTR-1.

**Figure 4.**
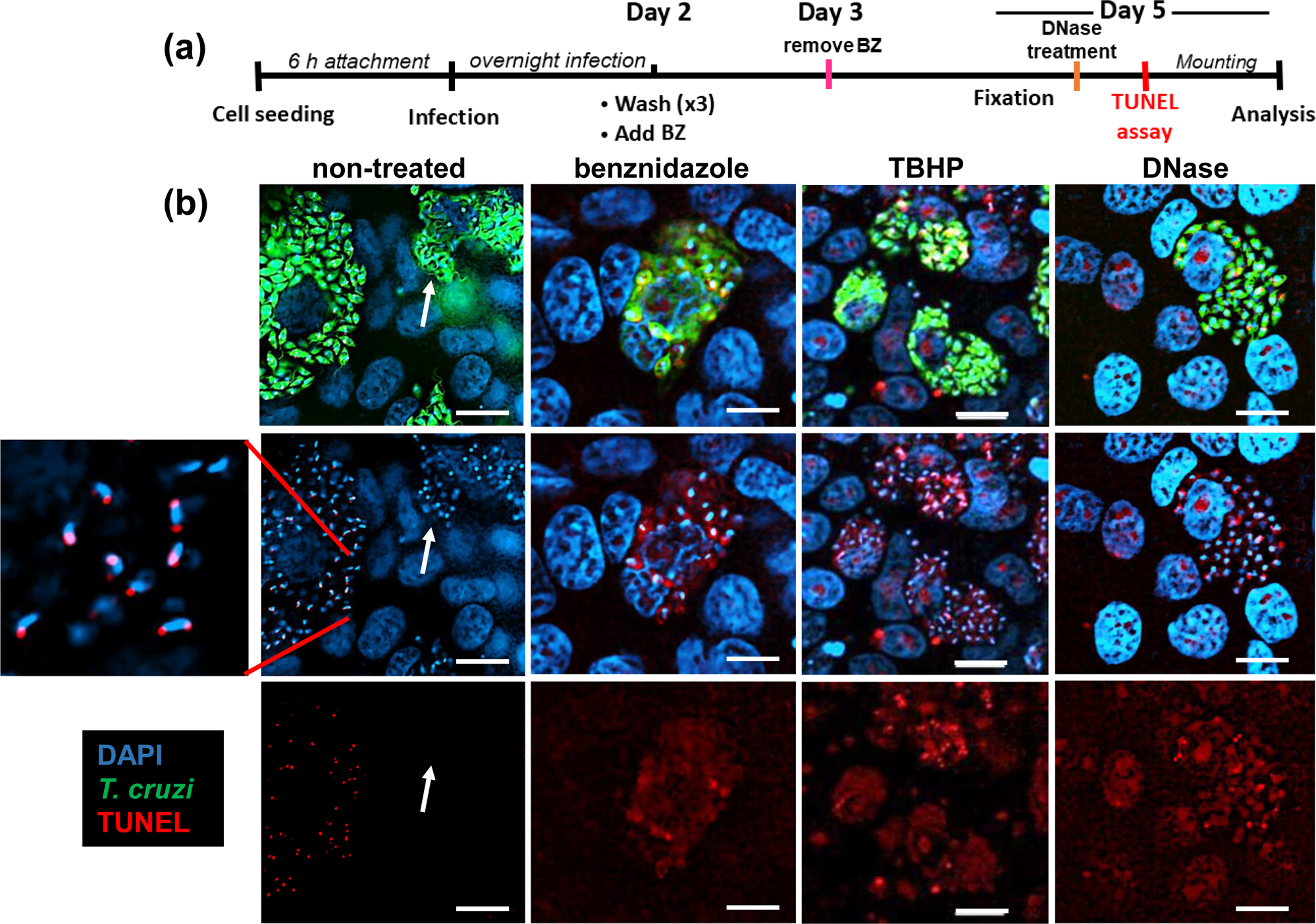
Fragmentation of *T. cruzi* DNA following benznidazole treatment. (a) Experimental outline. MA104 cells were infected with *T. cruzi* CL Luc::mNeon in 24-well plates. They were treated with either benznidazole (BZ) (200 µM) for 24 hours, or tert-butyl hydroperoxide (TBHP) (50 µM) for 3 days. Post-fixation, as a control group, untreated cells were treated with DNase. TUNEL assays were then performed and cells imaged using a Nikon Ti-2 E inverted microscope (Materials and Methods). (b) Representative images showing infected cells following each of the treatments. Parasites (green fluorescence), DNA (blue, DAPI), TUNEL (red). The enlarged inset (left) highlights replicating kinetoplast DNA (kDNA). The white arrows in the non-treated images show the location of a highly infected cell in which all the parasites have differentiated into TUNEL-negative non-replicating trypomastigotes. White scale bars = 10 µm.

### Tissue-specific survival of T. cruzi following benznidazole treatment

Mice in the pre-peak (day 9) and peak phases (day 14) of acute stage infections with the bioluminescent:fluorescent *T. cruzi* reporter strain CL-Luc::Neon [22] were treated once daily for 5 days with benznidazole at 25 mg/kg, a treatment regimen we had previously shown to be non-curative [33]. This resulted in a 97-99% knock-down in the whole-body parasite burden by the end of the treatment period, as inferred by *in vivo* bioluminescence imaging (Figure 5a). *Ex vivo* imaging was then used to examine organs and tissues. In non-treated mice, the infection was widely disseminated, with all organs and tissues highly bioluminescent (Figure 5b and c). Drug treatment resulted in a major reduction in both the parasite load and the number of infection sites, although complete parasite clearance was not observed. Importantly, there was no indication from the pattern of the remaining bioluminescent foci that any specific tissue or organ had acted as a location where parasites were preferentially protected from drug activity (Figure 5c). This is consistent with benznidazole pharmacokinetics in *T. cruzi* infected mice, where there is extensive bio-distribution of the drug amongst organs and tissues [34].

**Figure 5.**
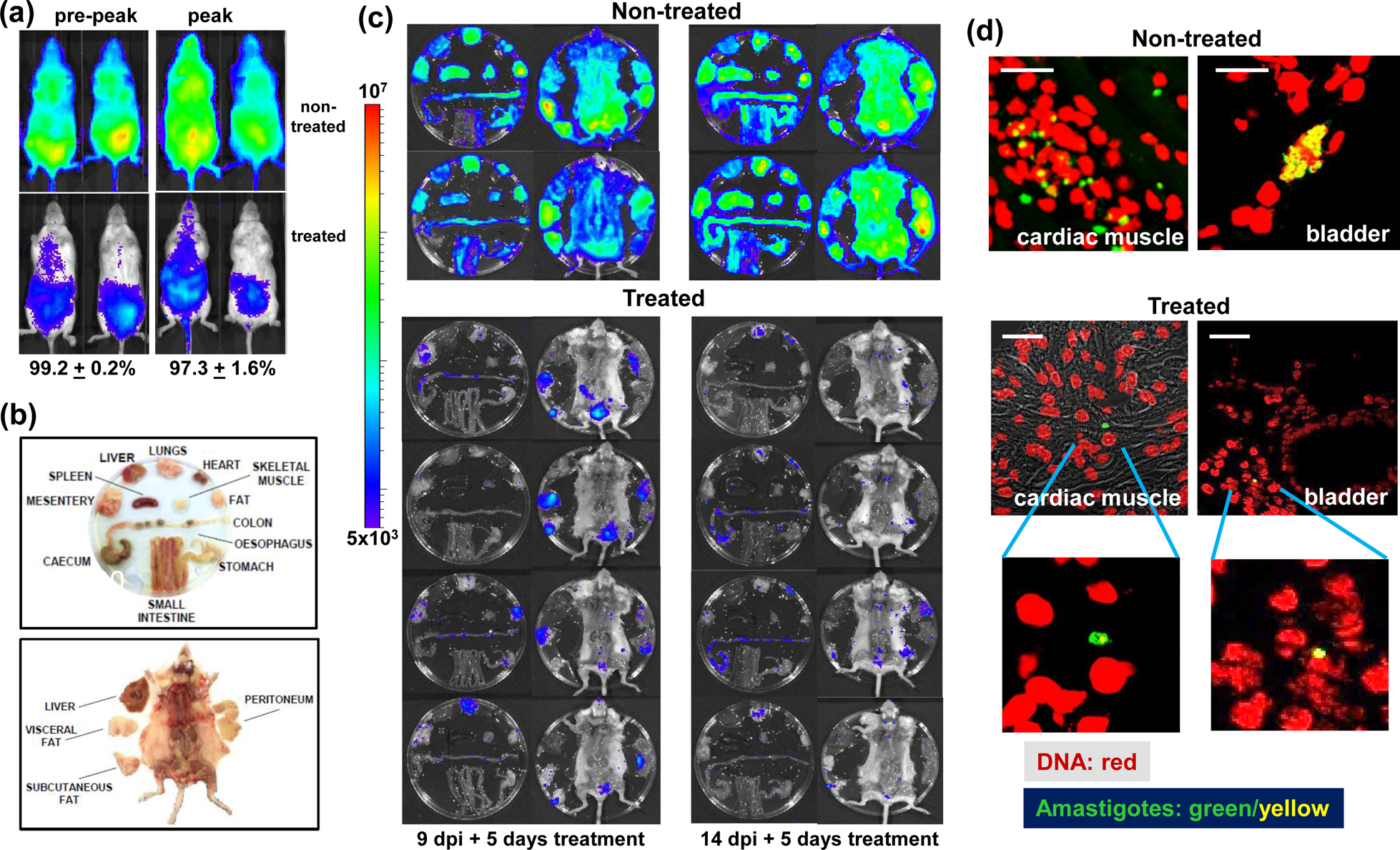
Monitoring the tissue-specific impact of non-curative benznidazole treatment by bioluminescence imaging. (a) Representative *in vivo* images of *T. cruzi* infected BALB/c mice after once daily oral treatment with 25 mg/kg benznidazole for 5 days (Materials and Methods). Treatment was initiated in the acute stage either 9 (pre-peak) or 14 (peak) days post-infection (dpi). The percentage drop in whole-body bioluminescence is indicated (n=6). (b) Schematic showing the arrangement of tissues, organs and carcass used for *ex vivo* imaging. (c) *Ex vivo* images of non-treated and treated infected mice. The heat-map for both *ex vivo* and *in vivo* imaging is on a log10 scale and indicates the intensity of bioluminescence from low (blue) to high (red) with minimum and maximum radiance values as indicated. (d) Fluorescent detection of parasites in the bladder and cardiac muscle of non-treated and treated mice during the acute stage of infection, using a Zeiss LSM880 confocal laser scanning microscope. DNA (red, DAPI); parasites (green fluorescence; yellow, if on a red background). White scale bars = 20 µm. Lower images show expanded view of single amastigote infections.

Tissue samples containing bioluminescent foci were excised and examined by confocal laser scanning microscopy (Materials and Methods). mNeonGreen fluorescence was detectable in all tissue samples obtained from infected non-treated mice in the pre-peak and peak phases of the acute stage, and parasite location could be established at single-cell resolution [25] (Figure 5d). In infected host cells, parasite numbers were then determined with precision using serial z-stacking (Figure 6a). Tissues examined included the heart, adipose tissue, bladder, spleen, peritoneum, lungs, liver, colon, rectum, cecum, stomach and skeletal muscle. Intracellular amastigote numbers varied considerably in non-treated mice. The vast majority of infected cells had a burden of less than 50 parasites, although occasional large “nests” that contained up to 150 parasites could be detected (1.5% of infected cells) (Figure 6b). Tissue and organs from benznidazole-treated mice were similarly processed on the day following treatment cessation. Infected cells were much less abundant, and were more difficult to locate. Across a range of tissue types, the majority of infected cells (>75%) contained only a single amastigote (Figure 6b). It is implicit therefore, that recrudescence after drug treatment does not arise from the survival of a small number of intact large nests. As with the situation *in vitro*, the most parsimonious explanation is that persisters are the survivors from intracellular populations where the other parasites have been killed. Alternatively, amastigotes in single/low-level infected cells may exist in a state where they are less susceptible to the trypanocidal activity of benznidazole. Treatment at higher doses (100 mg/kg, 5 days), or for longer periods (30 mg/kg, 10 days), also results in relapse (S1 Figure). However, immediately after treatment cessation with these regimens, it is more technically challenging to detect the small number of surviving tissue-resident parasites.

**Figure 6.**
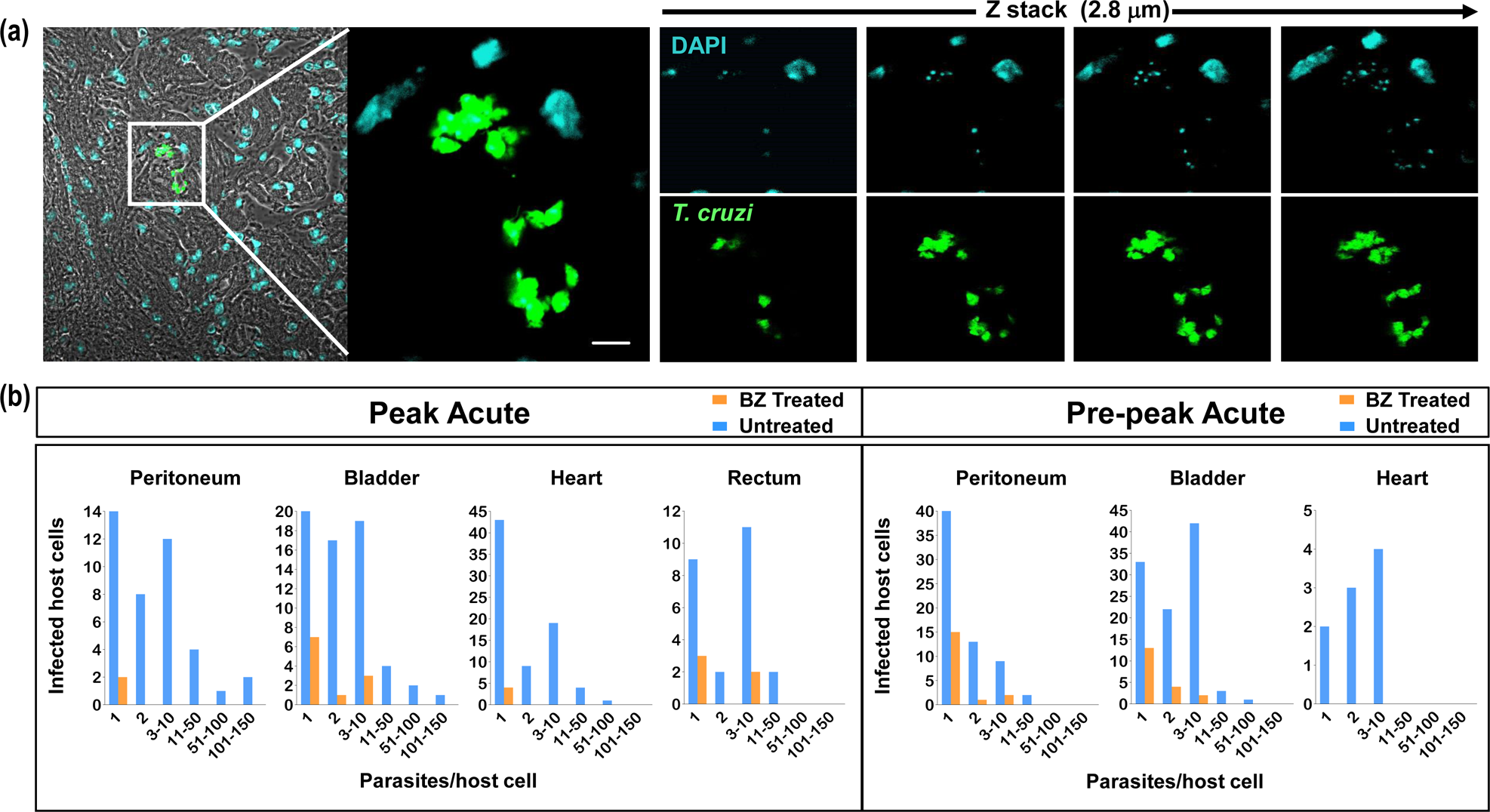
In benznidazole treated mice, the majority of cells that remain infected contain only a single parasite. BALB/c mice in the pre-peak (9 dpi) and peak (14 dpi) stages of infection were treated once daily with 25 mg/kg benznidazole for 5 days (as in Figure 5). Serial sections (10 µm) from a range of tissues were prepared and examined in 3-dimensions by z-stacking to determine the precise number of amastigotes in each infected cell (Materials and Methods) (see also S1 Video). (a) Illustrative 0.7 µm serial images across a section of cardiac tissue from a non-treated mouse. Amastigotes (green); DNA (blue, DAPI). Parasite numbers can be determined with precision by counting the distinctive intensely stained kinetoplast (mitochondrial) DNA that co-localises with green fluorescence across serial sections. White scale bar = 5 µm. (b) Parasite numbers per infected cell determined by exhaustive screening of multiple sections obtained from a range of tissues from treated (orange bars) and non-treated (blue bars) mice.

### Parasites that survive benznidazole treatment *in vivo* are in a non-replicative state

As above, BALB/c mice in the acute stage of infection were treated with a non-curative benznidazole dosing regimen (5 days, 25 mg/kg). Our strategy was then to use EdU incorporation into parasite DNA as a reporter [16,24], to determine the replicative status of intracellular amastigote persisters (Materials and Methods). However, at this point in the infection (day 20), many of the detected parasites displayed an irregular and diffuse morphology, in both treated and non-treated mice (S2a Figure, as example). In non-treated mice, this was associated with enhanced accumulation of proliferating host cells within infected cardiac tissue (cardiomyocytes are normally terminally differentiated) and a major increase in leukocyte infiltration (S2b Figure). We inferred from this that the observed parasite damage was mediated by the adaptive immune response. To avoid this confounding factor, which would complicate interpretation of the EdU incorporation data, we therefore switched murine models and used CB17 SCID mice, an immunodeficient strain that lacks functional lymphocytes [35]. This resolved the issue, and few proliferating cells were then observed in the cardiac sections (see Figure 7c, for one such example). At the analysis time-point, infected cardiac cells in untreated mice could be readily detected, with the vast majority containing between 1 and 50 morphologically intact parasites. Of the amastigotes surveyed, 45% (845/1863) were found to be in a replicative state as inferred from EdU incorporation (Figure 7a and b, S3 Figure, S1 Video). In benznidazole treated CB17 SCID mice, infected cardiac cells were much rarer, with a total of 23 infected cells detected after exhaustive searching of tissue sections. The majority of these contained only a single amastigote (74%, 17/23), and no host cells were found that contained more than 3 parasites (S3 Figure). None of the 30 persisting parasites detected in cardiac tissue after benznidazole treatment were in a replicative state, as judged by a lack of incorporation of EdU into genomic and/or kinetoplast DNA (Figure 7a and c, S2 and S3 Videos).

**Figure 7.**
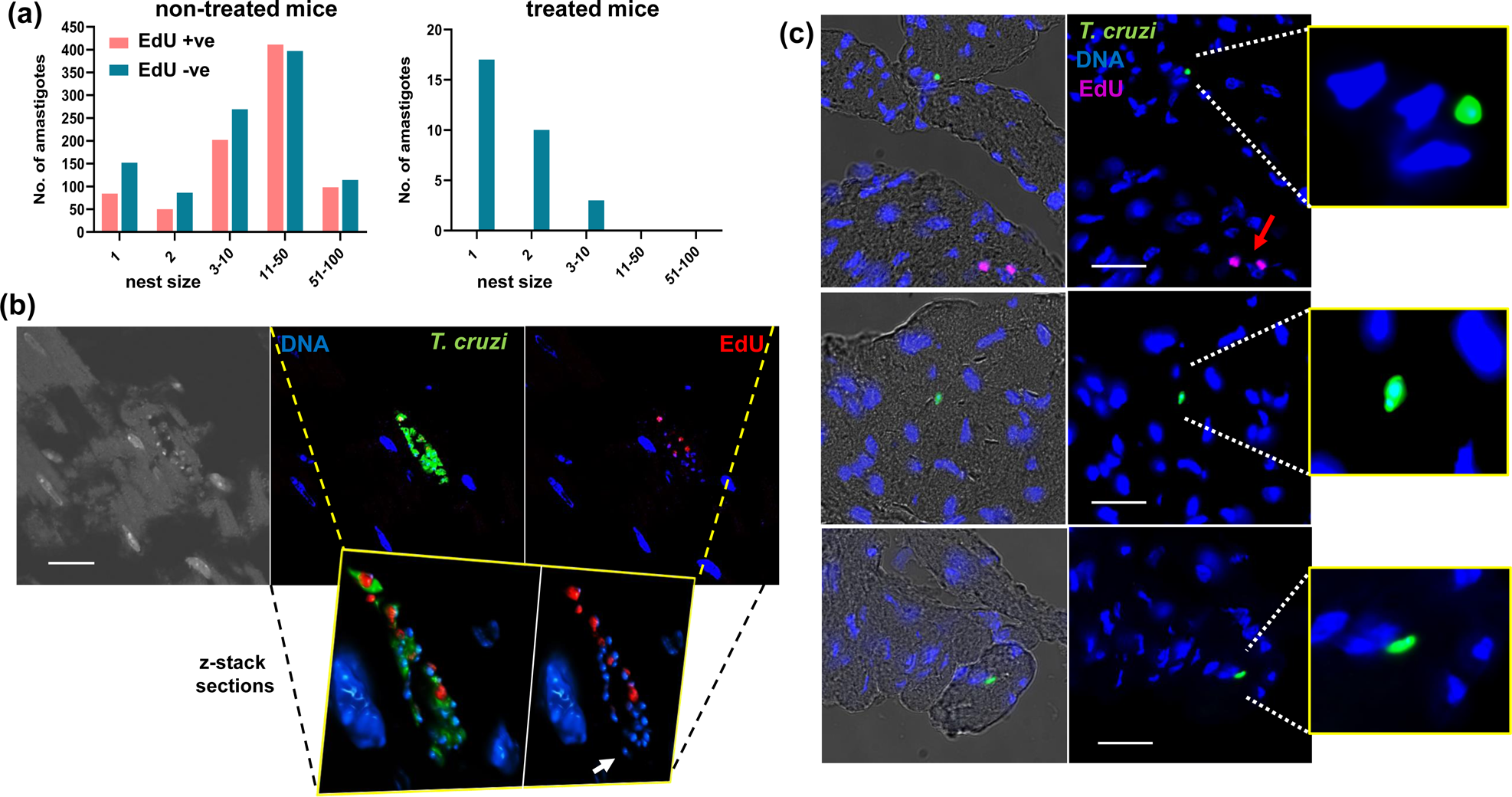
Parasites that persist in mouse cardiac tissue after benznidazole treatment are in a non-replicative state. CB17 SCID mice were infected with *T. cruzi* and 10 days post-infection they were treated once daily with 25 mg/kg benznidazole for 5 days. EdU labelling was then carried out as described previously (Materials and Methods). Mice were euthanised (16 dpi), and cardiac sections prepared and imaged using a Nikon Ti-2 E inverted microscope (9 mice per group, 3 randomly selected sections from each mouse) (a) Infection burden per infected cardiac cell (nest), with the number of EdU+ve/-ve parasites indicated (see also S3 Figure). (b) Images showing an infected cardiomyocyte from a non-treated mouse that contains both replicating and non-replicating amastigotes. The inset shows an example of a single serial cross section derived by 3-dimensional confocal laser scanning microscopy (z-stacking), which was used to determine the precise number of amastigotes in the infected cell (white arrow indicates intensely stained kinetoplast DNA) (see S1 Video for 3-dimensional image of this infected cell). DNA (blue, DAPI); EdU+ve amastigotes (red); parasites (green fluorescence). White scale bar = 10 µm. (c) Images of infected cardiomyocytes from benznidazole-treated mice. None of the amastigotes detected were EdU+ve. A purple arrow indicates two host cells that were in S-phase during the period of EdU exposure (upper image). White scale bars = 20 µM. The insets (right) show enlarged images of DAPI stained host cell nuclei and single infecting amastigotes (green). Full 3-dimensional images of parasites that persist after benznidazole treatment are shown in S2 and S3 Videos.

## Discussion

Benznidazole remains the front-line drug used to treat *T. cruzi* infections, despite multiple reports of treatment failure [9–11]. The underlying reasons for these non-curative outcomes have not been resolved. There is wide diversity in the levels of benznidazole sensitivity within natural *T. cruzi* populations [36]. However, this is not associated with polymorphisms in *TcNTR-1*, the gene that encodes the nitroreductase that initiates reductive activation of the drug [12]. Furthermore, inactivation of one *TcNTR-1* allele results in only a 2 to 4-fold increase in resistance [17], a level insufficient to account for the wide spectrum of benznidazole tolerance within natural populations (∼20-fold). Functional disruption of both *TcNTR-1* alleles confers ∼10-fold resistance, but this is associated with reduced infectivity that would prevent these highly-resistant parasites from becoming established within host populations [12,17]. Other mechanisms must therefore be involved [21].

In this study, we demonstrate that parasites which survive benznidazole treatment are not preferentially restricted to a specific organ or tissue in experimental mice (Figure 5c). This suggests that variable drug distribution is unlikely to be a major factor in recrudescence, at least in this context. This is in line with studies on benznidazole pharmacokinetics [34]. As an alternative explanation, it has been proposed that the replicative state of intracellular amastigotes could impact on drug sensitivity and that this might have a role in parasite persistence. This follows on from descriptions of apparent dormant forms of the parasite [14,37,38], stress-induced quiescence [15], and a reduced proliferation rate as an adaptation to chronic infection [16]. Here, we found that the small number of amastigotes that survive benznidazole treatment *in vitro* are in a non-replicative state (Figure 3). These persisters remain viable (Figure 2c), and at a population level, retain a capacity to replicate, differentiate and egress from host cells, even after 8 days exposure to 200 µM benznidazole (∼100x EC_50_) (Figure 1d). Within 9-11 days of treatment cessation, non-replicative parasites re-enter the cell cycle and to begin to proliferate (Figure 3).

Are these persister parasites derived from a pre-existing non-replicative sub-population, or do they enter this state as a response to some aspect of drug activity? In the case of benznidazole, damage to parasite DNA is a known mode of action [19,20,39–41]. Reductive drug metabolism is initiated by TcNTR-1, resulting in the production of reactive metabolites that ultimately break down to yield glyoxal [18,42]. These reactive molecules promote the formation of DNA crosslinks, mutagenesis, and chromosomal breaks. In addition, increased oxidative stress gives rise to the formation of oxidized nucleotides, such as 8-oxo-guanine. As further evidence for this mode of action, here we show that benznidazole treatment leads to the generation of extensive parasite-specific DNA breaks detectable by TUNEL assays (Figure 4). An inevitable consequence of this will be induction of the DNA damage response pathway [43], exit from the cell cycle, and recruitment and assembly of DNA repair enzymes at the sites of damage. Lesion repair, if successful, would then be followed by re-entry into the cell cycle and continued proliferation. Therefore, triggering of the *T. cruzi* repair pathways [44,45] by benznidazole may have the effect of inducing a transient non-replicative state that protects at least some parasites from further drug-mediated damage. This does not exclude mechanisms such spontaneous dormancy [14] or other forms of stress-induced quiescence [15] that could act in parallel, or in the case of drugs with a different mode of action, play a key role in persistence.

When the impact of benznidazole on murine infections was assessed, as with the situation *in vitro*, the few parasites that survived treatment were restricted mainly to host cells that contained only one or two amastigotes. This was the case across a wide range of tissue types (Figure 6 and 7), indicating that these scarce residual parasites are the likely source of recrudescence. To assess the replicative status of persisting amastigotes, we focused on cardiac tissue. The heart is the major site of pathology during both the acute (myocarditis) and chronic (cardiomyopathy) stages of Chagas disease. Results from the *in vivo* experiments closely mirrored those from *in vitro* studies. In non-treated mice, maximum nest size was typically up to 50 amastigotes, with 40-50% of the parasite population in S-phase (Figure 7a and b). In contrast, following treatment, nest size was much reduced (typically 1-2 parasites), and all detected parasites were non-replicative (Figure 7a and c, S2 and S3 Videos).

Remarkably, some parasites that are exposed to continuous benznidazole concentrations of 100xEC_50_ for 8 days *in vitro* are able to survive and proliferate (Fig. 1d). *In vivo*, drug exposure following a single dose (25 mg/kg) is considerably less than this, and plasma concentrations drop below the EC_50_ value within 12 hours of administration [33]. Nevertheless, as we show here, 5 days treatment of experimental mice at this dosing regimen is sufficient to kill the vast majority of parasites (Figure 5), with rare non-replicative amastigotes being the only survivors. Attempts to enhance benznidazole efficacy by modifying the dosing regimen have yielded differing results. In murine models, extended intermittent treatment (4 months, twice weekly administration) at 250 mg/kg yielded the best curative outcomes [46]. In contrast, with humans, the cure rate (∼80%) using the standard dose remained the same, irrespective of treatment length (once daily for 2 or 8 weeks) [47]. Benznidazole treatment protocols that overcome the problem of persister parasites may be possible, but extended regimens are likely to risk unacceptable toxicity, with a concomitant impact on patient compliance. Combination therapy could be one route to reducing treatment length and benznidazole dose, but clinical trials involving co-treatment with ergosterol biosynthesis inhibitors have shown little benefit [11,47]. New anti-*T. cruzi* drugs, delivered individually or in combination will require an ability to eliminate persisters. The recent report that cyanotriazoles can cure experimental infections through selective irreversible binding to topoisomerase II [48] highlights that covalent inhibition of essential enzymes could be one route to achieving parasite elimination.

## Materials and Methods

### In vitro parasite culturing

*T. cruzi* (CL Brener strain) that constitutively express a bioluminescent:fluorescent fusion protein (clone CL-Luc::Neon) were generated and cultured as described previously [22]. Metacyclic trypomastigotes were produced by transfer of epimastigotes to Graces-IH medium and harvested after 4-7 days. Tissue culture trypomastigotes (TCTs) were generated after infecting MA104 cells (an African green monkey kidney epithelial cell line) with metacyclic trypomastigotes [16].

### Sorting of benznidazole-treated T. cruzi infected cells

T25 tissue culture flasks were seeded with 10^6^ MA104 cells, incubated for 6 hours to allow attachment, and then infected with TCTs at an MOI of 10:1 (parasite:host cell). 18 hours later, external parasites were removed by thorough washing (×3), fresh supplemented Minimum Essential Medium Eagle (MEM, Sigma-Aldrich) was added, and benznidazole (Epichem Ltd.) made to a final concentration of 20 µM. The plates were incubated at 37°C, with fresh medium/benznidazole renewed on day 4. After 8 days, the cultures were washed (x2) and cells detached by incubation of the monolayers in TrypLE express (Thermo Fisher). For staining nuclei of live cells, monolayers were incubated with Hoechst 3342 (200 ng/ml) for 90 minutes prior to detachment. To differentiate viable/non-viable cells, cellular suspensions were incubated with propidium iodide (PI) (1 µg/ml) for 30 minutes. Following staining and fractionation, the cell suspension was sorted under CL3 conditions using an Aria BD Cell Sorter, with the laser setting appropriate for the stained/fluorescent infected cells.

### TUNEL assays

Mammalian cell cultures infected with *T. cruzi* were grown on coverslips in 24-well plates as described above. At specific time points, monolayers were fixed with 4% paraformaldehyde in PBS, air-dried, washed (x1) in PBS, permeabilized in 0.1% TritonX-100/PBS for 5 minutes, and washed (x3) with PBS. 20 µl TUNEL reaction mixture (*In situ* Cell Death Detection Kit, TMR-red, Roche) was then added to each coverslip and the reaction incubated in the dark for 1 hour at 37°C. The coverslips were washed (x3) in PBS, mounted on slides with VECTASHIELD® with DAPI (Vector Laboratories, Inc.), and then examined using Zeiss LSM880 confocal or inverted Nikon Ti-2 E inverted microscopes.

### In vitro labelling with EdU

As a marker for DNA replication, infected cells were labelled with 5-ethynyl-2′-deoxyuridine (EdU). At various time points after the cessation of benznidazole treatment, fresh medium containing 10 µM EdU (Sigma-Aldrich) was added to selected wells [16] and cultures incubated for a further 6 hours. Monolayers were then washed (x2) and incubated for 45 minutes in 4% paraformaldehyde diluted in PBS. Finally, coverslips were removed and washed in PBS (x2). The extent of EdU incorporation was determined using a Click-iT PlusEdU AlexaFluor 555 Imaging kit (Invitrogen), followed by washing with PBS (x2), and the coverslips then mounted in VECTASHIELD® with DAPI. Cells were imaged in three dimensions with a Zeiss LSM880 confocal microscope.

### Ethics statement

Animal work was performed under UK Home Office project licences (PPL 70/8207 and PPL P9AEE04E4) and approved by the LSHTM Animal Welfare and Ethical Review Board. Procedures were performed in accordance with the UK Animals (Scientific Procedures) Act 1986.

### Murine infections

CB17 SCID mice and BALB/c mice were purchased from Charles River (UK). Animals were maintained under specific pathogen-free conditions in individually ventilated cages, with a 12 hour light/dark cycle, and access to food and water *ad libitum*. Female CB17 SCID mice, aged 9 weeks, were infected with 1×10^3^ TCTs in 0.2 ml 10% fetal calf serum in Dulbecco Minimal Essential Medium (DMEM), with 4.5g/litre glucose, via i.p. injection. BALB/c female mice, aged 7-8 weeks, were infected by i.p. injection of 1×10^3^ BTs derived from CB17 SCID mouse blood. At experimental end-points, mice were culled by exsanguination under terminal anaesthesia.

For drug treatment, benznidazole was prepared for administration at 2.5 or 10 mg/ml in 5% dimethyl sulfoxide (v/v)/95% HPMC suspension vehicle (0.5% (w/v) hydroxypropyl methylcellulose, 0.5% (v/v) benzyl alcohol, 0.4% (v/v) Tween 80 in Milli-Q water). Mice were treated under the regimens outlined in the legends to the relevant figures, with the drug administered by oral gavage [33]. Non-treated control mice were administered with 0.2 ml 5% dimethyl sulfoxide (v/v)/95% HPMC suspension vehicle.

### Bioluminescence imaging

Infected mice were injected with 150 mg/kg d-luciferin i.p., then anaesthetized using 2.5% (v/v) gaseous isoflurane 5-10 minutes after d-luciferin administration [49,50]. They were then placed in an IVIS Lumina II Spectrum system (PerkinElmer) and ventral and dorsal images acquired using Living Image v4.7.3. Exposure times varied between 30 seconds and 5 minutes, depending on the signal intensity, and anaesthesia was maintained throughout via individual nose cones. For *ex vivo* imaging, mice were injected with d-luciferin, and euthanised as above, then perfused via the heart with 10 ml 0.3 mg/ml d-luciferin in PBS. Organs and tissues were removed and transferred to a Petri dish in a standardized arrangement, soaked in 0.3 mg/ml d-luciferin in PBS, and imaged using maximum detection settings (5 minutes exposure, large binning). The remaining animal parts and carcass were checked for residual bioluminescent foci, also using maximum detection settings. The detection threshold for *in vivo* imaging was determined using uninfected mice.

### Histological procedures

To ensure preservation of fluorescence in tissue samples derived from infected mice, we adapted methodology previously described [51]. Bioluminescent foci identified by *ex vivo* imaging were excised from tissue and fixed in pre-chilled 95% ethanol for 20-24 hours at 4°C in histology cassettes. Samples were dehydrated (4×15 minute washes with 100% ethanol), cleared (2×12 minute washes with xylene) and embedded in paraffin wax. Tissue sections (3-10 µM) were cut with a microtome. For confocal imaging, slides were melted on a heat pad for 30 minutes, further de-paraffinized with two changes (12 minutes each) of xylene, three changes (12 minutes each) of Tris-buffered saline, pH 7.6 (TBS), permeabilised with 0.1% Triton X-100 + 0.1% sodium citrate, and then mounted using VECTASHIELD® with DAPI, before storing at 4°C in the dark until required. For immunostaining, slides were blocked and stained with a 1:250 dilution of rat anti-mouse CD45 primary antibody (Tonbo Biosciences; clone 30-F11, cat. #70-0451-U100), used in combination with a 1:500 secondary Alexa Fluor 647 donkey anti-rat antibody (Invitrogen Thermo Fisher Scientific, cat. #A-21209).

### Confocal and wide-field fluorescence microscopy

For imaging, we used a Zeiss LSM880 confocal laser scanning microscope, with the Zen black software. Accurate determination of intracellular parasite numbers was carried out by 3-dimensional imaging (z-stacking), with the appropriate scan zoom setting [25]. Mounted slides were also imaged using a Nikon Ti-2 E inverted microscope, with images processed using Zen blue software for analysis. Samples were imaged using a Plan Apo 60x oil immersion objective (NA = 1.42, Ph2, Nikon) and an ORCA Flash 4.0 CMOS camera (Hamamatsu). For each specimen, parasites were detected using the green fluorescent channel. In regions of interest, a z-stack of 30 to 50 images with an axial spacing of 0.3 µm was taken for a series of fields of view along the length of the tissue to account for the 3D location of the parasite within individual cells. 3D-video projections were acquired and processed using the NIS Advanced Research software package.

### In vivo labelling with EdU

To identify host cells and parasites undergoing DNA replication, mice were given 2 EdU i.p. injections (12.5 mg/kg, 6 hours apart) in PBS at the specific time points as detailed in the Results section. The mice were then left overnight, euthanised by terminal anaesthesia, and tissue sections fixed and sectioned as above. Labelling of incorporated EdU was carried out using the Click-iT Plus EdU AlexaFluor 555 Imaging kit as described for cultured monolayers.

## Acknowledgements

This work was supported by UK Medical Research Council (MRC) grants MR/T015969/1 to J.M.K. and MR/R021430/1 to M.D.L., and funding from the Drugs for Neglected Diseases initiative (DNDi). DNDi received financial support from: Department for International Development (DFID), UK; Federal Ministry of Education and Research (BMBF) through KfW, Germany; and Médecins sans Frontières (MSF) International. AIW was in receipt of an MRC LID (DTP) Studentship (MR/N013638/1).

The funders had no role in study design, data collection and analysis, decision to publish, or preparation of the manuscript.

