## Supplementary information text for "*Trypanosoma cruzi* persisters that survive benznidazole treatment *in vitro* and *in vivo* are in a transient non-replicative state"

**S1 Figure. *In vivo* imaging of BALB/c mice in the acute stage of *T. cruzi* infection after non-curative benznidazole treatment.** (a) Dorsal images of mice treated orally, once daily, with 30 mg/kg benznidazole for 10 days (Materials and Methods). (b) Ventral images of mice treated orally, once daily, with 100 mg/kg benznidazole for 5 days. In both instances, treatment was initiated 14 days post-infection (dpi) (indicated with red arrows). Heat-maps are on log10 scales and indicate the intensity of bioluminescence from low (blue) to high (red), with minimum and maximum radiance values as indicated.

**S2 Figure. Leukocyte infiltration into cardiac tissue of a BALB/c mouse 20 days post-infection with *Trypanosoma cruzi.*** Female BALB/c mice aged 6-8 weeks were infected with *T. cruzi* CL Luc::mNeon (Materials and Methods). 19 dpi, the mice were given two EdU i.p. inoculations (12.5 mg/kg) 6 hours apart. They were then left overnight, euthanised, and tissue sections prepared and imaged by fluorescence microscopy (Materials and Methods). (a) Intracellular parasites in cardiac tissue imaged using a Zeiss LSM880 confocal laser scanning microscope. Note the irregular and diffuse parasite morphology. DNA (blue, DAPI); nuclei of EdU+ve host cells (turquoise); parasites (green fluorescence). White scale bar = 5 µm. (b) Sections of cardiac tissue from uninfected and *T. cruzi* infected mice showing infiltration of CD45+ cells (red), imaged using a Nikon Ti-2 E inverted microscope. A yellow 200 µm diameter circle is shown for reference.

**S3 Figure. The replicative status of intracellular *T. cruzi* parasites in cardiac sections from benznidazole treated and non-treated CB17 SCID mice.** Mice were infected with *T. cruzi* and 10 days later treated, or not, with 5 daily doses of benznidazole (25 mg/kg). Mice were then inoculated with EdU (two doses of 12.5 mg/kg, 6 hours apart) (Materials and Methods), culled the following day, and 3 randomly selected cardiac sections from each mouse (n=9, per group) were searched exhaustively for infected cells containing fluorescently labelled parasites. Images were acquired using a Nikon Ti-2 E inverted microscope. The numbers of EdU+ve parasites in each group are shown.

**S1 Video.** **3-dimensional imaging of a *T. cruzi* infected cardiomyocyte in a tissue section from a non-benznidazole treated CB17 SCID mouse**. See legend to S3 Figure for further details. DNA (blue, DAPI); parasites (green fluorescence); EdU+ve amastigote DNA (red).

**S2 Video. 3-dimensional imaging of an infected cardiomyocyte containing a single amastigote persister in a tissue section from a benznidazole treated CB17 SCID mouse.** See legend to S3 Figure for further details. DNA (blue, DAPI); parasites (green fluorescence). This tissue section was also assessed for EdU incorporation into DNA (red), but no parasite or host cell EdU positivity was detected. In the upper image, the dimensions of the infected host cell are indicated by a dashed yellow line.

**S3 Video. 3-dimensional imaging of an infected cardiomyocyte containing two amastigote persisters in a tissue section from a benznidazole treated CB17 SCID mouse.** See legend to S3 Figure for further details. DNA (blue, DAPI); parasites (green fluorescence). This tissue section was also assessed for EdU incorporation into DNA (red), but no parasite or host cell EdU positivity was detected. Two amastigotes are present in the infected cell, with their kDNA visible as a discrete blue disc.
