## Supplementary Figures 1 - 3 for "*Trypanosoma cruzi* persisters that survive benznidazole treatment *in vitro* and *in vivo* are in a transient non-replicative state"

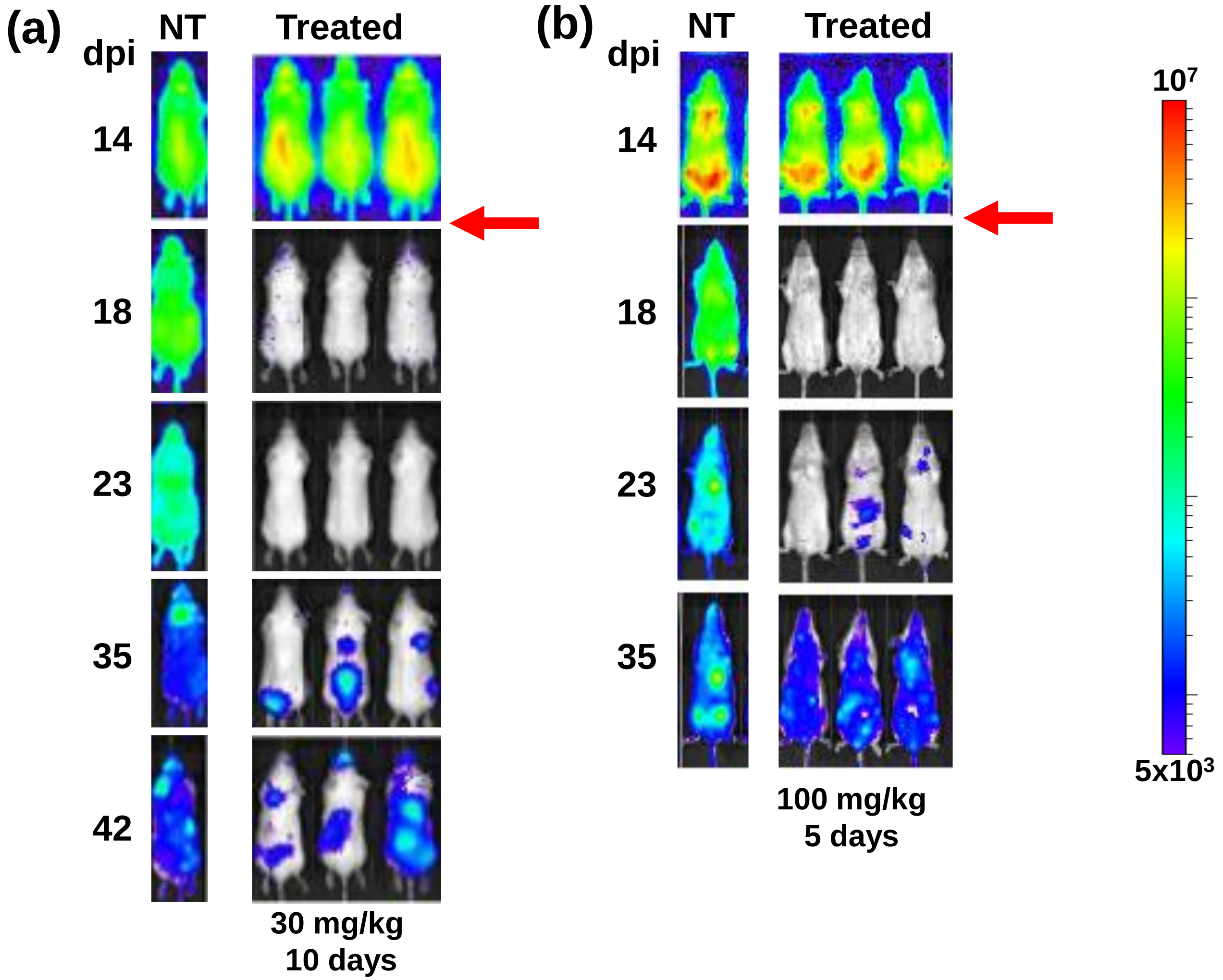
(a)
dpi
NT
Treated
14
18
23
35
42
30 mg/kg
10 days
(b)
dpi
NT
Treated
14
18
23
35
100 mg/kg
5 days
$10^7$
$5x10^3$

S1 Fig.

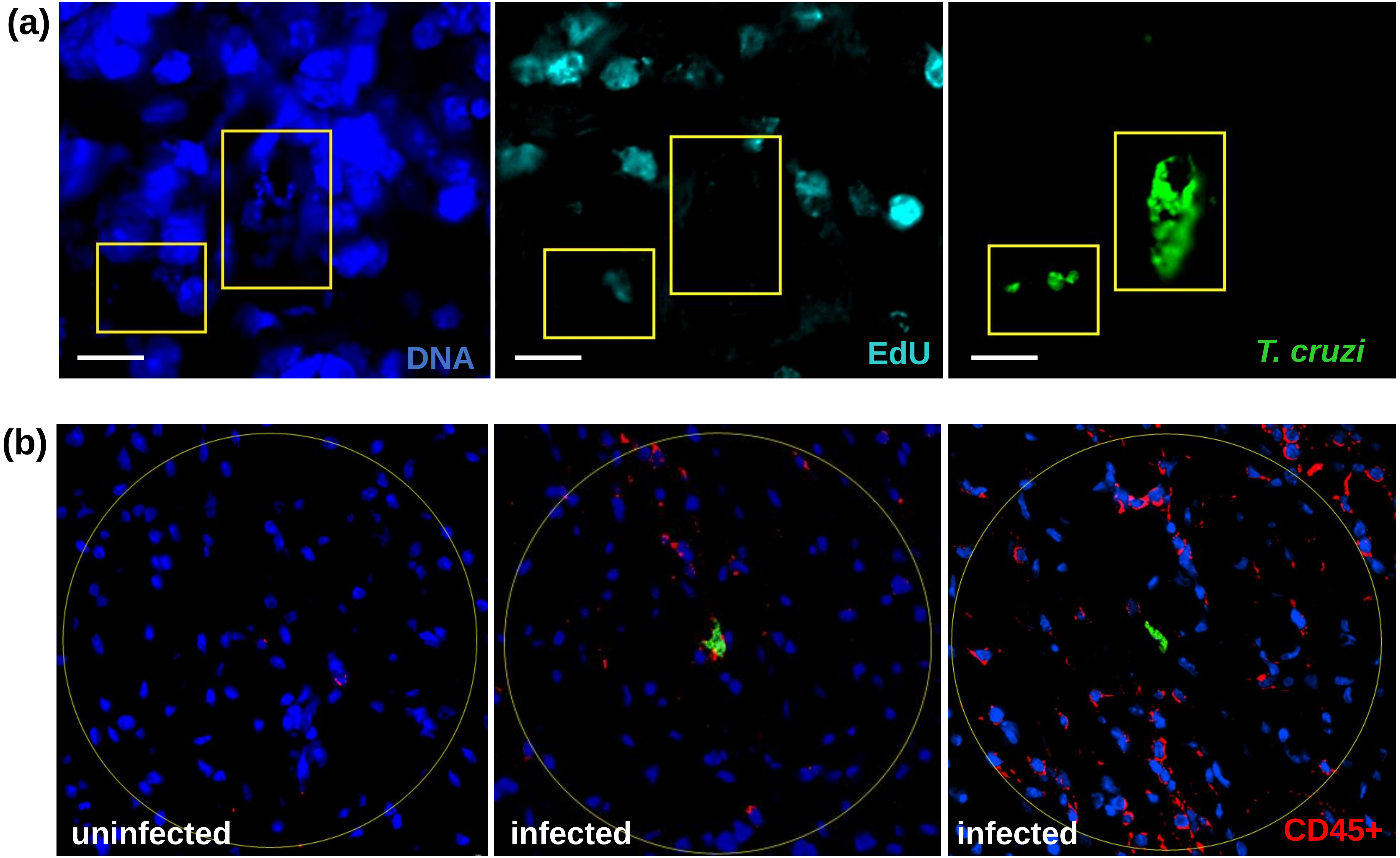


S2 Fig.

| ACUTE STAGE, VEHICLE TREATED SCIDS | | | | |
|---|---|---|---|---|
| Nest size | # infected cells | # amastigotes | # EdU+ve amastigotes | # EdU-ve amastigotes |
| 1 | 236 | 236 | 84 (36%) | 152 (64%) |
| 2 | 68 | 136 | 50 (37%) | 86 (63%) |
| 3-10 | 84 | 471 | 202 (43%) | 269 (57%) |
| 11-50 | 45 | 808 | 411 (49%) | 397 (51%) |
| 51-100 | 4 | 212 | 98 (46%) | 114 (54%) |

| ACUTE STAGE, BENZNIDAZOLE TREATED SCIDS | | | | |
|---|---|---|---|---|
| Nest size | # infected cells | # amastigotes | # EdU+ve amastigotes | # EdU-ve amastigotes |
| 1 | 17 | 17 | 0 | 17 (100%) |
| 2 | 5 | 10 | 0 | 10 (100%) |
| 3-10 | 1 | 3 | 0 | 3 (100%) |
| 11-50 | 0 | 0 | 0 | 0 |
| 51-100 | 0 | 0 | 0 | 0 |

S3 Fig.
